# Transcriptional Changes Fade Prior to Long-Term Memory for Sensitization of the *Aplysia* Siphon-Withdrawal Reflex

**DOI:** 10.64898/2025.12.28.696784

**Authors:** Tania Rosiles, Melissa Nguyen, RJ Calin-Jageman, IE Calin-Jageman

## Abstract

Forming a long-term memory requires changes in neuronal transcription. What happens, though, as the memory is forgotten? And how does the transcriptional state relate to the maintenance and recall of the long-term memory? To answer these questions we have been systematically tracing the time-course of transcriptional changes evoked by long-term sensitization in the marine mollusk *Aplysia californica*. Our approach captures transcriptional changes in neurons of known behavioral relevance using a within-subjects design, delineating patterns of transcriptional change that are comprehensive and reproducible. We have previously reported that within 1 day of long-term sensitization training there is a widespread transcriptional response involving robust changes in over 5% of tested transcripts (1,252 of ∼22k; Conte, 2017). Within 1 week, however, memory strength fades and nearly all transcriptional changes relapse to baseline (Perez, 2018). Here we report microarray analysis (N = 16) of transcriptional changes 5 days post-learning, a time-point when memory strength has weakened but is still robust. Remarkably, we find that at this intermediate behavioral stage nearly all transcriptional changes have fully decayed, even in subsets of animals that have shown very little forgetting. Thus, most transcriptional changes seem to decay more rapidly than memory expression. We discuss several possible ways that memory expression could become decoupled from detectable transcriptional regulation.

**Highlights:**

- Long-term sensitization training produces a memory that then fades over the course of a week, with behavioral expression at 5 days at an intermediate stage of partly forgotten with continued clear sensitization and considerable variety across animals.
- The transcriptional response to sensitization training fades more quickly than behavioral expression, with nearly all transcripts regulated 1 day after training showing a statistically significant decline in regulation, even amongst animals that had shown little forgetting.
- Transcription does not seem to have a straightforward relationship with the expression of sensitization memory, with a small set of transcripts consistently regulated even as behavioral expression changes and strong behavioral expression possible without most of the transcriptional changes observed during early maintenance.

## 1. Transcriptional Changes Fade Prior to Long-Term Memory for Sensitization of the *Aplysia* Siphon-Withdrawal Reflex

Long-term memories are distinguished not only by their duration but also by their requirement for changes in neuronal gene expression (Goelet et al., 1986). After a memory is induced, though, how do transcriptional states relate to memory expression? The “consensus model” is that transcriptional changes trigger the encoding of long-term memory (Klann & Sweatt, 2008) and that this includes activation of self-sustaining biochemical changes that perpetuate at least some transcriptional changes to support the ongoing expression of the memory (e.g. Smolen et al., 2019; Zhang et al., 2010). Under this model, forgetting is a disruption of maintenance mechanisms, and transcriptional states and memory expression should be tightly linked. There is some evidence, though, that the relationship between transcription and memory expression is not straightforward. For example, some transcriptional changes seem to persist even after memory expression weakens (Kim et al., 2012; Perez et al., 2018), and savings memory (the rapid re-learning of a seemingly forgotten memory) can be expressed without notable changes in gene expression (Rosiles et al., 2020). A critical goal, then, is better characterizing the ways in which changes in neuronal gene expression sculpt not only the induction but also the maintenance and forgetting of long-term memory.

We have been studying this issue by tracing the changes in gene expression that accompany long-term sensitization of the tail-elicited siphon-withdrawal reflex in *Aplysia californica* (Figure 1A). Sensitization is an evolutionarily conserved form of memory (Walters, 2018) in which a painful stimulus causes an increase in reflex responsiveness, often accompanied by additional behavioral changes and modulated by the current motivational state of the animal (Leod et al., 2018; Shields-Johnson et al., 2012). In *Aplysia*, long-term sensitization can be induced through repeated noxious shocks applied to one side of the body (Scholz & Byrne, 1987; Wainwright et al., 2002), and then observed as a long-term, unilateral increase in the duration of the tail-elicited siphon-withdrawal, a defensive reflex (Walters & Erickson, 1986). The circuitry mediating this reflex has been characterized, and long-term sensitization has been shown to be due, in part, to long-term increases in the excitability and synaptic strength of the VC nociceptors located in the pleural ganglia (Scholz & Byrne, 1987; Walters et al., 2004). This form of learning has proven especially useful for studying transcriptional correlates of long term memory because a) the unilateral expression of sensitization enables powerful within-subjects comparisons of gene expression, and b) analysis of gene expression can proceed from the pleural ganglia which contain the VC nociceptors, providing an excellent transcriptional signal from neurons of known behavioral relevance to the induction and maintenance of sensitization memory (Calin-Jageman et al., 2024).

**Figure 1:**
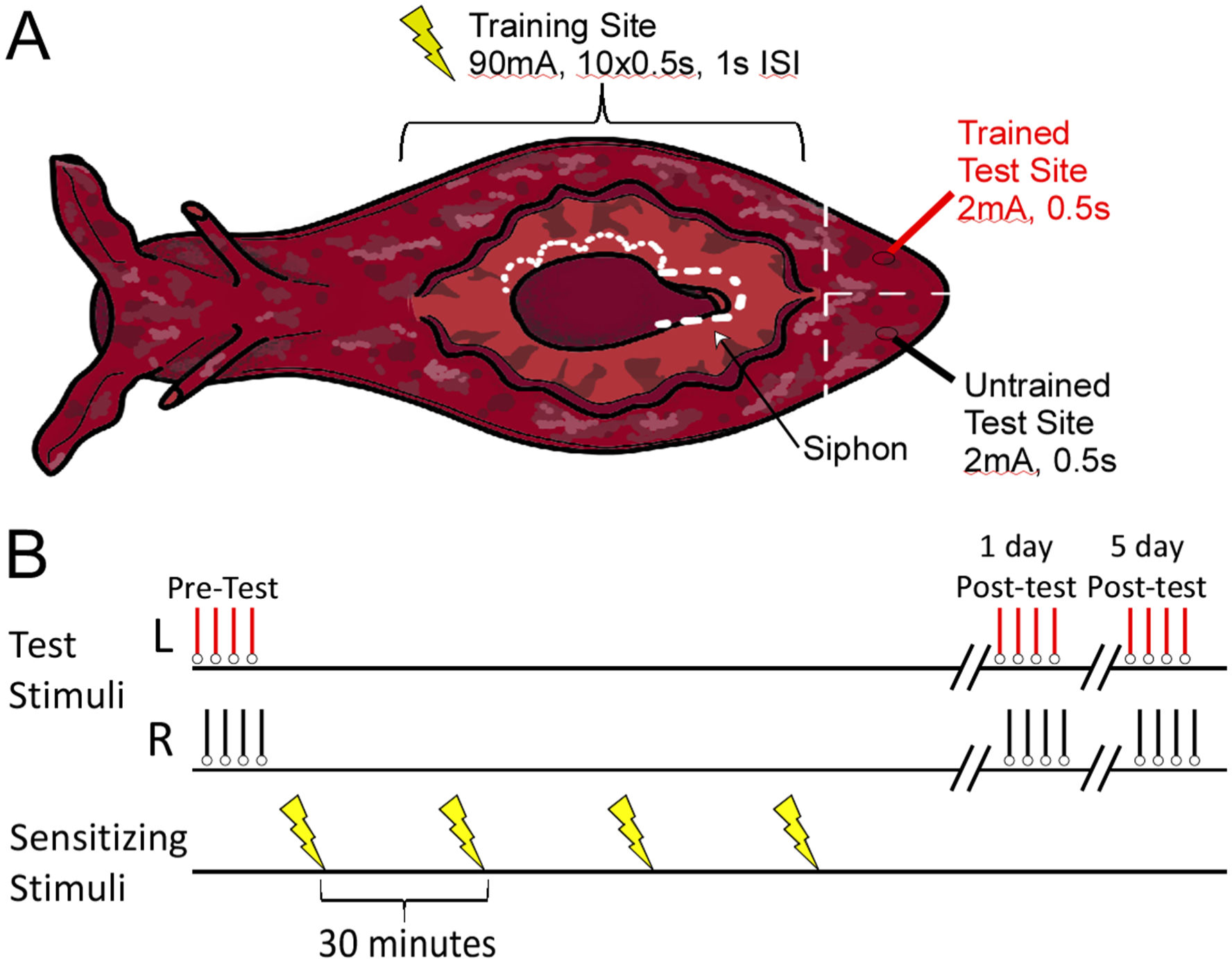
Long-erm sensitization of the tail-elicited siphon-withdrawal reflex (T-SWR). (A) Cartoon diagram of the body of an Aplysia. T-SWRs are evoked by applying an innocuous shock to the left or right tail (arrows). The duration of the T-SWR serves as an index of behavioral responsiveness. For LTS training, a noxious shock is applied along the length of one side of the body (lightning bolts). (B) Experimental protocol. First, baseline T-SWR measures are made on the left and right side of the tail, then LTS training is applied to one side of the body, then T-SWR measures are made again 1 and 5 days after post tests. Immediately after the 5-day post tests, pleural ganglia from the trained and untrained side are harvested in matched pairs of left-trained and right-trained animals.

We have previously shown that long-term sensitization training produces widespread changes in gene expression in the pleural ganglia containing VC nociceptors (as well as directly in the VC neurons themselves). One hour after training, there is notable upregulation of about 80 transcripts, many of which are transcription factors (Herdegen, Conte, et al., 2014; Herdegen, Holmes, et al., 2014). At 1 day after training, this transcriptional response ramifies, involving over 1,200 transcripts that show clear regulation (Conte et al., 2017). After 1 week, the expression of sensitization fades completely (reflex responsiveness returns to baseline), and so to do most transcriptional changes, with the vast majority of transcripts regulated 1-day after training showing a statistically significant decline in regulation (Perez et al., 2018). Although *most* transcriptional decay within 1 week we have identified 7 transcripts which show “beyond forgetting” regulation that persists for up to 2 weeks, long after recall of sensitization has fully decayed (Patel et al., 2018; Perez et al., 2018). Interestingly, sensitization memory can be rapidly relearned (savings or latent memory; Philips et al., 2006) and can then re-persist in a long-term form (for more than 1 day), but this does *not* re-activate transcriptional changes (Rosiles et al., 2020), providing a puzzling disconnect between transcriptional states and memory expression.

Here we characterize the transcriptional changes occurring 5 days after long-term sensitization training, a time-point at which there is substantial forgetting and yet still clear expression of long-term sensitization memory. Following a pre-registered design, we designed our study to help answer 4 questions:

- What is the fate of the ∼1200 transcripts strongly regulated during early maintenance of long-terms sensitization?
- Is the small set of “beyond forgetting” transcripts also regulated at this earlier time point?
- Does the passage of time and forgetting of sensitization produce additional, late-breaking transcriptional changes?
- Do any gene expression changes predict individual levels of forgetting?

## 2. Materials and Methods

Prior to conducting microarray analysis we pre-registered our sample size plan, data exclusions, all manipulation and measures, and a complete microarray analysis script (https://osf.io/g3ueu). We then followed our pre-registered plan exactly, with no notable deviations. We report below all pre-registered analyses and note as exploratory any follow-up analyses developed after data collection. Our preregistration documents, analysis scripts, and behavioral data are posted to the Open Science Framework (https://osf.io/ccwks/); the microarray data is also posted to NCBI’s Gene Expression Omnibus (Geo ID: forthcoming).

### 2.1. Animals

Animals (75-125g) were obtained from the RSMAS National Resource for *Aplysia* (Miami, FL) and maintained at 16° C in one of two 90-gallon aquariums with continuously circulating artificial sea water (Instant Ocean, Aquarium Systems Inc.). Handling was as described previously (Herdegen, Holmes, et al., 2014). *Aplysia* are true hermaphrodites.

### 2.2. Long-term sensitization training

A one-day long-term sensitization training protocol (Figure 1B) was used, adapted from Wainwright et al. (2002) but with a stronger shock (90mA vs. 60mA) and a constant-current stimulus (Bonnick et al., 2012; Cyriac et al., 2013). Training consisted of 4 rounds of noxious shock applied at 30 minute intervals to one side of the body with a hand-held electrode. Each round of shock consisted of 10 pulses (60Hz biphasic) of 500ms duration at a rate of 1hz and an amplitude of 90mA. During the course of each shock, the stimulating electrode was slowly moved from anterior (just behind neck) to posterior (just in front of tail) and back to cover nearly the entire surface of that side of the body. Side of training was counterbalanced.

### 2.3. Behavioral measurement

As a behavioral outcome, we measured the duration of the tail-elicited siphon-withdrawal reflex (T-SWR; see Walters & Erickson, 1986). The reflex was evoked by applying a weak shock to one side of the tail using a hand-held stimulator (60Hz biphasic DC pulse for 500ms at 2ma of constant current). T-SWR behavior was measured as the duration of withdrawal from the moment of stimulation to the first sign of siphon relaxation. Behavioral measurements were made by a researcher blind to the side of training.

To track sensitization memory, we measured T-SWR durations before training (Baseline) and then 1 and 5 days after long-term sensitization training (Post-Tests).

We focused on changes in T-SWR behavior by calculating the log-fold change from Baseline to Post-Test at each timepoint on both the trained and untrained side of each animal:

*LFC* = Log_2_(*Post-Test* / *Baseline*).

By this metric a score of 0 represents no change in T-SWR duration, and scores above 0 represent sensitization. We then calculated the degree of sensitization expression at each time point by comparing the change in T-SWR behavior on the trained side to the untrained side:

*Sensitization_Expression* = *LFC*_Trained_ - *LFC*_Untrained_

Finally, we calculated a forgetting index as the change in sensitization expression from day 1 to day 5:

*Forgetting_Score* = *Sensitization_Expression*_Day1_ – *Sensitization_Expression*_Day5_

With this metric, a score of 0 occurs when memory expression is stable (same expression on day 1 and day 5 leads to a forgetting score of 0), and is above 0 (indexing forgetting) when memory expression is lower on Day 5 than on Day 1.

### 2.4. Isolation and processing of pleural ganglia RNA

We compared gene expression in the pleural ganglia on the trained vs. untrained sides. To control for lateralized gene expression, samples from two animals trained on opposite sides were pooled (*N* = 32 animals) to create 16 sets of trained vs. control microarray samples.

To analyze transcription, pleural ganglia RNA was isolated immediately after the 5-day post test. These ganglia contain the VC nociceptors which help mediate the expression of long-term sensitization memory through training-induced long-lasting increases in excitability and synaptic efficacy (Cleary et al., 1998). Isolation and homogenization was exactly as described in Herdegen et al. (2014).

### 2.5. Sample size determination

We set a target of 16 microarray samples (requiring 32 animals). Our previous work has shown good sensitivity with 8 microarray samples; we doubled this target to account for the fact that regulation 5 days after training might grow more subtle than observed 1 day after training. Our sample size target exceeds the consensus recommendation of at least 5 biological replicates per group (Allison et al., 2006; Pavlidis et al., 2003; Tsai et al., 2003).

### 2.6. Quality controls

To ensure suitable samples, several quality controls were utilized to select animals for microarray analysis:

- Animals had to exhibit strong learning, defined as at least a 30% increase in T-SWR duration from baseline to the 1-day post-test.
- The expression of sensitization had to be unilateral, with less than a 30% increase in T-SWR on the untrained side from baseline to the 1-day post-test.
- There could be no protocol errors in training, testing, or isolation that would yield ambiguity about the processed samples (e.g a training shock applied to wrong side)

We ran animals in batches of 12-30. We completed training of 77 animals. Of these, 7 were excluded for exhibiting a weak training response, 6 were excluded for bilateral expression of sensitization, and 31 were excluded due to protocol errors (3 due to a single shock applied to the wrong side during training; 29 due to poor or uneven RNA isolation). This left 32 animals which were paired into 16 samples for microarray analysis. To pair animals for a microarray sample we first sorted qualifying animals within a batch by forgetting score and then combined two animals with similar forgetting scores but opposite sides of training (the left-trained animal with highest forgetting score was paired with the right-trained animal with highest forgetting score, and so on).

### 2.7. Microarray processing

We used the Aplysia Tellabs Array (ATA: GEO: GPL18666) to characterize changes in gene expression due to long-term sensitization training. This array includes 26,149 distinct probes representing all known sources of *Aplysia californica* ESTs and mRNAs at the time of design (January 2012). Based on estimates from previous microarray designs (Moroz et al., 2006), the ATA should cover >50-60% of all transcripts epressed in the nervous system. Full details on the array design are reported in Herdegen et al. (2014).

Microarray processing was completed by Mogene Inc. (St. Louis, MO). A two-color approach was used with each array hybridized to a sample from a trained or untrained animal. In half of cases, trained samples were hybridized with Cy3 and controls to Cy5; the other half we dye-swapped. Processing was exactly as described in Herdegen et al. (2014).

We identify microarray probes by the NCBI accession number for the EST or mRNA that the probe was designed to. For EST-based probes, when there is a definitive match of the EST to a gene model in the current *Aplysia* genome we also provide that accession number.

### 2.8. Statistical analysis

Behavioral responses were averaged by time point. Paired comparisons were made from baseline to post-test for each side. Standardized effect size estimates (cohen’s *d*) are corrected for bias (Hedges, 1981) and calculated so that positive values represent an increase in response (sensitization).

Microarray data was analyzed using limma (Ritchie et al., 2015a; Smyth, 2005) from the Bioconductor suite of tools (Gentleman et al., 2004) for R (Ihaka & Gentleman, 1996). Our processing script for identifying differentially regulated transcripts was pre-registered and is posted on the Open Science Framework. Median expression values were analyzed (Zahurak et al., 2007). These were corrected for background using the normexp+offset algorithm recommended for Agilent microarrays by Ritchie et al. (Ritchie et al., 2007). An offset of 30 was selected based on inspection of MA Plots. Expression was then normalized using the loess function (Smyth & Speed, 2003). Where multiple probes were used to measure the same EST or mRNA, these were averaged. Finally, trained and control expression were compared using an empirical Bayes-moderated *t*-test (Smyth, 2004). Statistical significance was calculated using Benjamini-Hochberg correction for multiple comparisons to maintain a 5% overall false-discovery rate (Benjamini & Hochberg, 1995). We used the treat function from limma (McCarthy & Smyth, 2009) to conduct a stringent test for significant regulation. Specifically, rather than use a null hypothesis of no regulation, we tested for regulation statistically distinguishable from at least a 10% change in expression in either direction. We have previously found that using this type of high-stringency criterion yields very strong predictive validity in independent qPCR (Herdegen, Holmes, et al., 2014; Holmes et al., 2014). We also quantified the degree of relationship between the regulation observed 5 days after LTS training with our previous screen of regulation observed 1 day after LTS training (Herdegen, Holmes, et al., 2014) and 7 days after LTS training (Perez et al., 2018). We examined the correlation between transcriptional states after correcting expression scores for potential measurement error using the genuine association of gene-expression profiles function (genas) in limma (Ritchie et al., 2015b). Finally, we conducted exploratory analysis of the completion of each gene list using the propTrueNull function (Ritchie et al., 2015b) and the convex decreasing densities approach developed by Langaas et al. (2005).

## 3. Results

### 3.1. Long-Term Sensitization Training Produces Robust Unilateral Sensitization That Is Partly Forgotten Within 5 days

All animals (*N* = 32 qualified animals) received long-term sensitization training (Figure 1B). This produced a robust sensitization memory on the side of training (Figure 2), with responses increasing from an average of 11.1 s at baseline to an average of 20s at the 1-day post test, nearly a doubling in response time (*LFC*_Trained_ = 0.84 95% CI [0.78, .91]). As expected, sensitization was unilateral, as on the untrained side response durations were at 11.2s at baseline and 11.1s at the 1-day post test, a negligible change in response (*LFC*_Untrained_ = 0.0 95% CI [-0.06, 0.04]). Thus, sensitization memory was strongly expressed 1 day after training: *Sensitization_Expression*_day1_ = *LFC*_Trained_ – *LFC*_Untrained_ = 0.85 95% CI[0.76, 0.94], *d*_avg_ = 5.2 95% CI [4.2, 6.5]).

**Figure 2:**
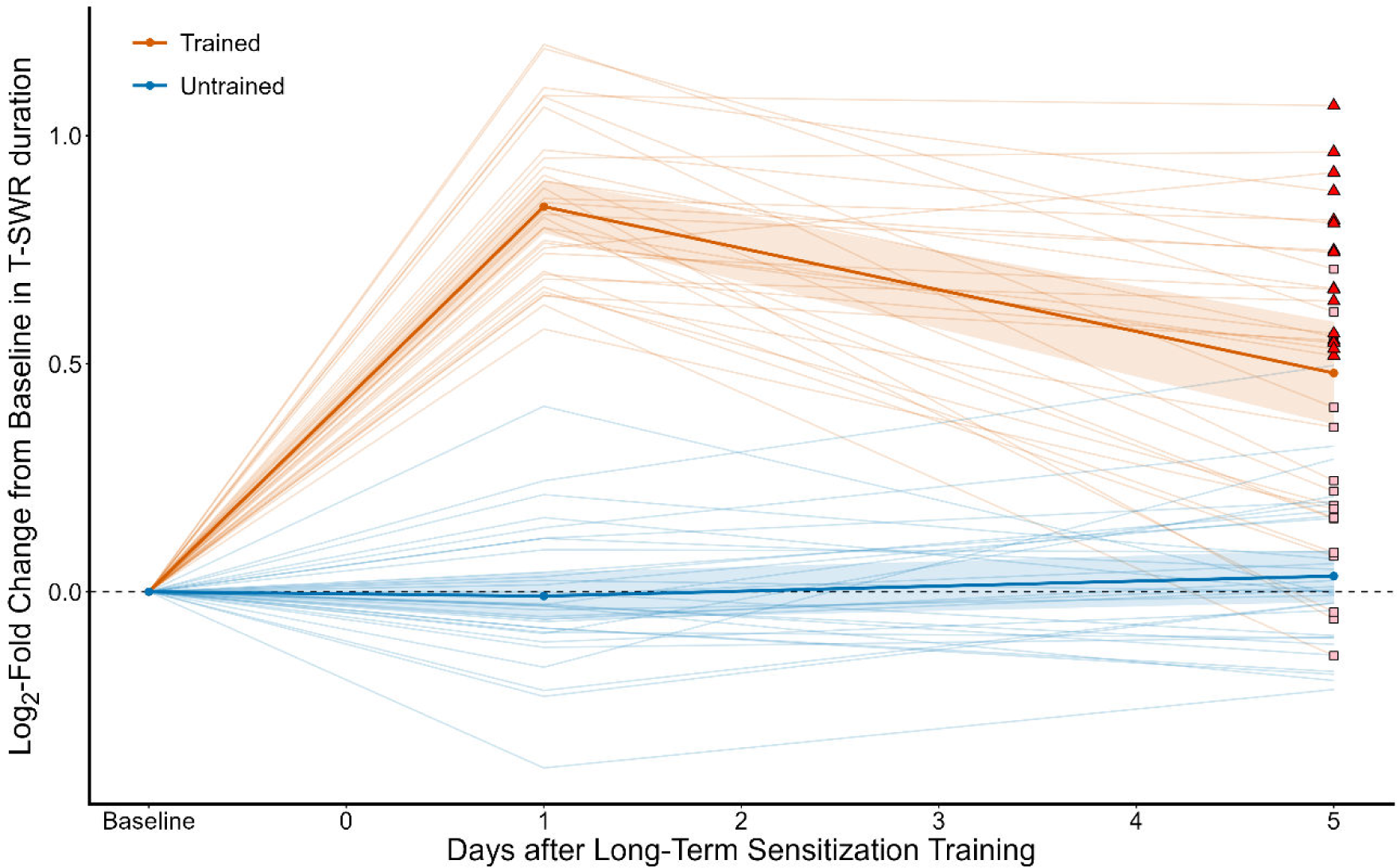
Training produces a long-term sensitization Memory that is partly forgotten at 5 days. This figure shows T-SWR duration as the log-fold change from baseline on both the trained (red) and untrained (black) sides. Dark lines with dots represent group means; shading indicates 95% CI of the mean. Individual animals are represented by the light lines, with a symbol for the 5-day post test indicating if it was grouped into the low-forgetting group (red triangles)or high-forgetting group (pink squares) for an exploratory analysis to determine if there was more prominent gene expression changes in animals in the low-forgetting animals.

Five days after training sensitization was substantially reduced but still clearly expressed. On the trained side, responses declined to a mean of 16.0s at the 5-day post test, only a 50% increase from baseline rather than the doubling observed 1-day after training (*LFC*_Trained_ = 0.48 95% CI [0.36, 0.60]). As expected, responses on the untrained side were stable, with a mean of 11.4s, very similar to baseline measures (*LFC*_Untrained_ = 0.03 95% CI [-0.02, 0.09]). Thus, there was still substantial expression of sensitization on day 5 (*Sensitization_Expression*_day5_ = 0.45 95% CI[0.30, 0.59], *d*_avg_ = 1.7 95% CI [1.1, 2.4]), but with notable forgetting relative to Day 1 (*Forgetting_Score* = *Sensitization_Expression*_day1_ - *Sensitization_Expression*_day5_ = 0.41 95% CI [0.29, 0.53], *d*_avg_ = = 1.2 95% CI [0.8, 1.6]). The mean forgetting score represents a loss of about ½ of the initial expression of sensitization, but there was considerable diversity in forgetting scores, with some animals showing almost no change in expression and others showing essentially no further sensitization.

Overall, our protocol succeeded in capturing a partially forgotten state of sensitization where there is clear and yet notably weaker expression of sensitization, with considerable diversity between animals. To capture this diversity, we matched each left-trained animal with a right-trained animal with a similar forgetting score, and we used the average forgetting score for each microarray sample in several analyses reported below.

### 3.2. To What Extent is Regulation 1 Day After Sensitization Preserved 5 Days After Training?

How similar is gene regulation 5 days after training to the pattern observed 1 day after training? To answer this question, we compared changes in gene expression across these time points, focusing specifically on 1,252 transcripts which we have previously identified as strongly regulated 1 day after training (Conte et al., 2017). Surprisingly, we found that regulation at day 5 is only modestly correlated with regulation at day 1 (*r* = .38 95% CI [.33, .43], *r*_corrected_ = .54, *N* = 1,252, Figure 3a). Moreover, the slope of the relationship was weak (*B* = 0.07 95% CI [0.06, 0.08]), indicating that, on average, regulation 5 days after training is less than 10% as strong as what was observed 1 day after training. For comparison, we have previously conducted the same analysis (Rosiles et al., 2020) on two independent samples both collected 1 day after training and found a very strong correlation in expression patterns (*r* = .95 95% CI[.93, .96], *r*_corrected_ = .99).

**Figure 3:**
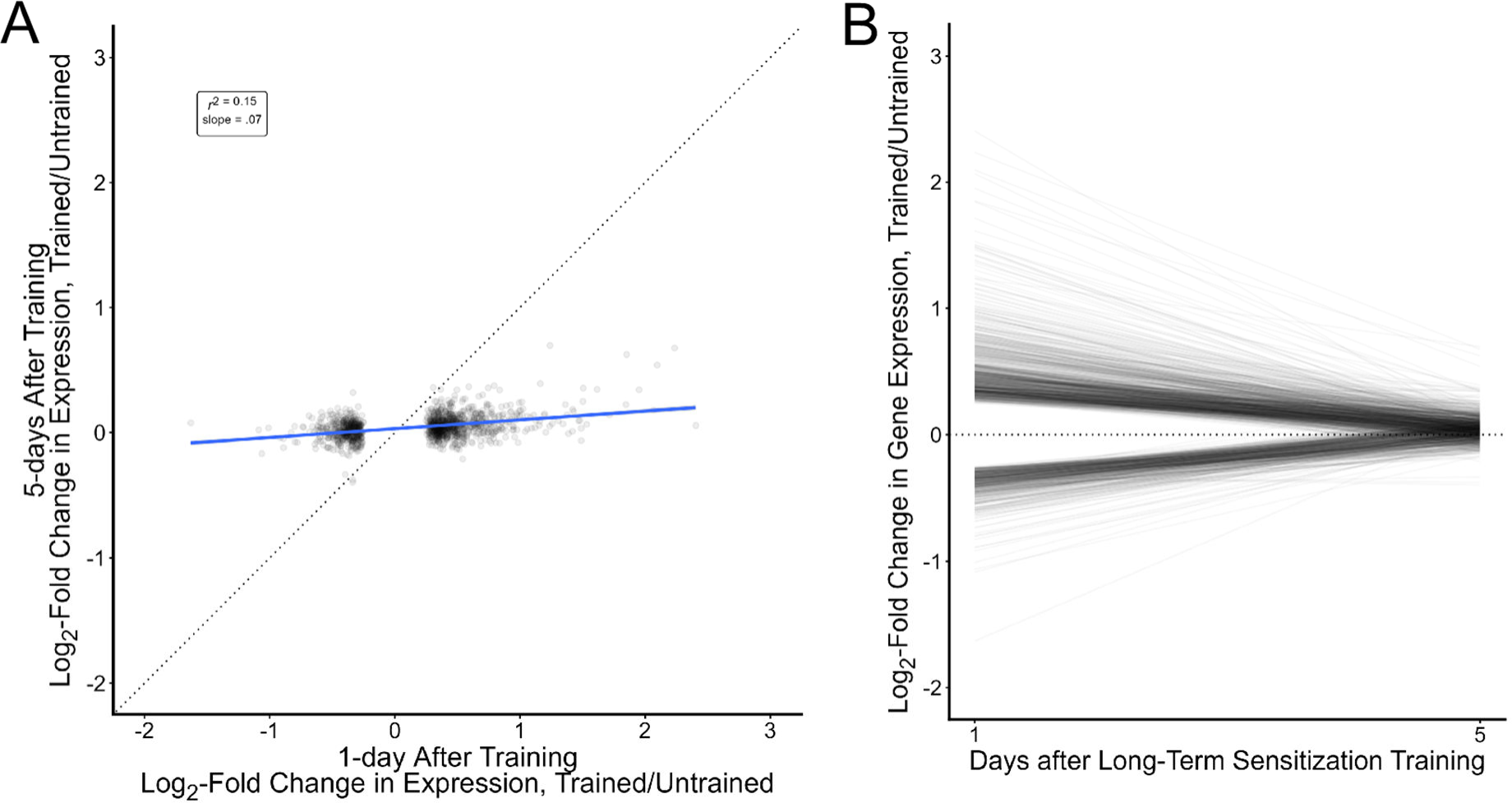
Gene expression changes observed 1 day after training have mostly decayed by 5 days after training. A) Scatterplot of log-fold changes in gene expression (Trained vs. Untrained) 1 and 5 days after long-term sensitization data for each of the 1,252 transcripts significantly regulated 1 day after training. The dashed line has a slope of 1 indicating the relationship expected if regulation was perfectly preserved over time. The blue line represents the actual line of best fit and its 95% confidence interval (shading). The large difference between the unit line and actual slope indicates a substantial attenuation of learning-induced gene expression changes. B) The same data but each line representing regulation from day 1 (starting point) to day 5 (ending point) for a transcript. The dashed line at 0 indicates no regulation. The 1-day data in this figure is from Conte et al., 2017.

Consistent with the correlational analysis, when we tested for significant regulation in the current data set, we found that *none* of the 1,253 transcripts regulated 1 day after training still showed clear regulation 5 days after training (none showed a statistically significant difference between trained and untrained expression against a null of 10% or less change in expression). With 16 paired samples and a focused analysis, this is probably not due to a lack of power. Indeed, analysis of the distribution of p values suggested a false-positive rate of 0.0001, giving an expected value of 0 false negatives over these 1,253 comparisons. Figure 3B shows the time-course for each of these transcripts, with a line representing regulation at day 1 (from Conte et al., 20) to day 5 (current experiment); the sharp decline towards no regulation notable in nearly all transcripts indicates the substantial attenuation of regulation that has occurred as time has elapsed since training.

We followed up these pre-registered analyses with three exploratory analyses. First, we directly compared regulation at days 1 and 5. Consistent with the correlational analysis, this showed that 94% (1,181) of the transcripts previously identified as regulated at day 1 showed a statistically significant decline in regulation at day 5; this suggests that the lack of regulated transcripts is not due simply to poor power, but to a statistically reliable decline in regulation for most transcripts. We also explored using a less-stringent criteria for identifying regulated genes (a null of 0 rather than of at least 10%).

This increased the number of transcripts qualifying as regulated from 0 to 39 (Supplemental Table 1), still a very small proportion (3%) of the transcripts which had been clearly regulated 1 day after training. Finally, we isolated our analysis to only the samples of animals which had shown low levels of forgetting. Specifically, we split the 16 microarray samples into a “high forgetting” set and a “low forgetting” set (see Figure 2, the triangle and square symbols denote the animals in these constructed groups). As expected for intentionally dividing the samples in this way, this produced an enormous difference in forgetting scores across these post-hoc groups (*M*_High_forgetting_ = 0.67 95% CI [0.56, 0.79], *M*_Low_forgetting_ = 0.14 95% CI [0.07, 0.22], *d*_avg_ = 2.8, 95% CI [1.8, 3.9]), with the low-forgetting group having lost, on average, only 17% of their initial expression of sensitization. We then tested for significant regulation of expression only in the low-forgetting group. This did not substantially alter conclusions: the correlation with day 1 regulation was actually slightly weaker among the low-forgetting samples (*r* = .35 95% CI [.30, .40], *r*_corrected_ = .50, *N* = 1,252) and the number of transcripts passing a stringent test for regulation remained at 0.

Overall, these results suggest that most of the gene regulation apparent 1 day after training substantially weakens within 5 days, though some of the pattern of regulation persists, and at low stringency a small number of transcriptional changes are detectable. This was true even in animals which did not show substantial forgetting of sensitization.

### 3.3. To What Extent Are Transcripts Regulated for 7 Days Similarly Regulated at 5 days?

We have previously shown that a small set of 7 transcripts remain regulated after forgetting, with some showing clear learning-associated changes in expression for up to 2 weeks after training (Patel et al., 2018; Perez et al., 2018). We next examined the extent to which these transcripts are regulated 5 days after training (Figure 4).

**Figure 4:**
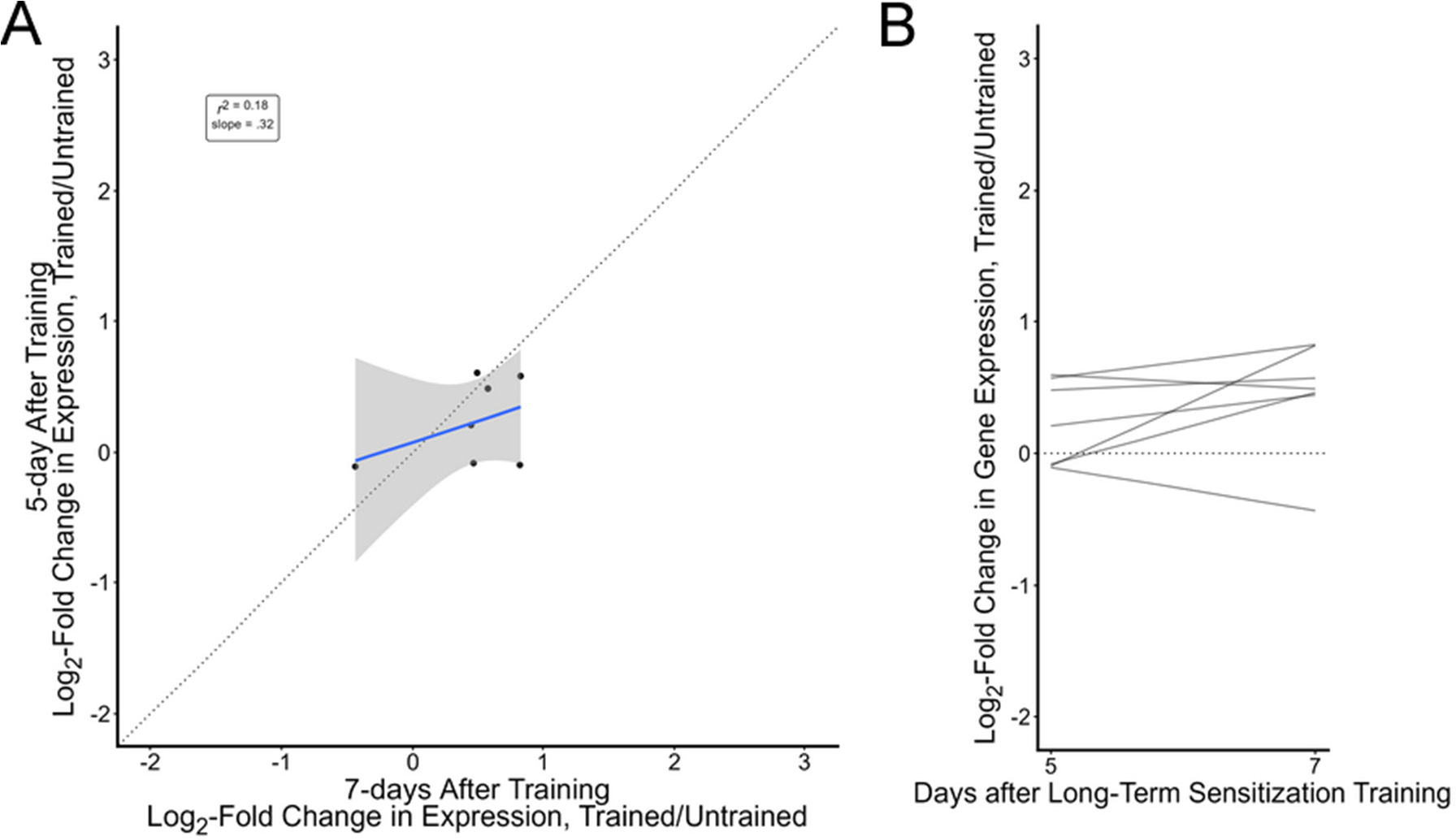
The small core of transcripts regulated 7 days after training are mostly detectable 5 days after training. A) Scatterplot of log-fold changes in gene expression (Trained vs. Untrained) 5 and 7 days after long-term sensitization data for the 7 transcripts significantly regulated 7 days after training. The dashed line has a slope of 1 indicating the relationship expected if regulation was perfectly preserved over time. The blue line represents the actual line of best fit and its 95% confidence interval (shading). B) The same data but each line representing regulation from day 5 (starting point) to day 7 (ending point) for a transcript. The dashed line at 0 indicates no regulation. The 7-day data in this figure is from Perez et al., 2018.

We found that 4 of these very-persistently regulated transcripts also showed clear regulation at this earlier timepoint (AH005259.2 FRMFamide precursor; FF066943.1/XM_013087893.2; EB254334.1; EB257711.1/XM_013081020.2). In an exploratory analysis with a less-stringent test this increased to 5 (EB255259.1/XM_035971949.1 spectrin). Two transcripts expected to be up-regulated were instead very slightly down-regulated in this sample (EB243511.1/XM_005100516.3 ankyrin repeat and BTB/POZ domain-containing protein 1; EB342172.1). Overall, levels of regulation in these transcripts were correlated with what we had observed 1 week after training (*r* = .42 95% CI[-.48, .89], *r*_corrected_ = .71, *N* = 7, Figure 4A), though the small sample size makes this relationship highly uncertain. Given that we have previously demonstrated that these transcripts are very persistently regulated after training, these results were not surprising. These analyses demonstrate, though, that the study design and approach is sensitive enough to detect a high proportion (5 of 7) of regulated transcripts.

### 3.4. To What Extent Are New Transcriptional Changes Induced During Forgetting?

We next examined if the passage of time might activate additional transcriptional changes that we have not yet observed as related to sensitization, analyzing the 24,839 microarray probes not included in the previous two analyses. After correction for multiple comparisons, no transcripts were flagged as regulated. Although this is a large number of tests for 16 paired samples, the lack of detected regulation did not seem due to low power, as the estimated false-positive rate was 0 at this stringency level.

We followed up this pre-registered analysis with an exploratory analysis using a lower level of stringency (null of 0). This identified 47 transcripts which could potentially be late-regulated (Supplemental Table 2). Of these, 22 showed a statistically significant difference in regulation from what had been observed at day 1. Even at this low level of stringency, the estimated false-positive rate remained at 0. Thus, at low-stringency there is modest evidence for some late-breaking transcriptional changes, but with a high risk of flagging what may be negligible levels of regulation.

### 3.5. To What Extent Are Gene Expression Changes Related to Forgetting?

We examined if changes in gene expression might predict forgetting, assessing the correlation between gene expression and forgetting scores across the entire microarray. After correction for multiple corrections, we found no transcripts showing a statistically significant correlation with forgetting scores.

As an exploratory analysis we increased power by restricting the analysis just to the 1,259 “day 1” transcripts (fewer comparisons requires less aggressive correction, increasing power). This restricted analysis, however, also failed to identify any transcripts significantly correlated with forgetting scores. Similarly, testing the set of 7 persistently regulated transcripts also failed to identify any that showed a significant association with forgetting scores.

A sample size of 16 is small for a correlational analysis, especially with correction for multiple comparisons. However, the main issue was likely the lack of gene regulation observable at this time point (restriction of range).

## 4. Discussion

We examined the transcriptional correlates of a partly forgotten memory. We found that transcription decays more rapidly than behavior, with very little of the pattern of transcriptional regulation observed 1 day after training preserved 5 days after training; this was true even in a subset of animals which had shown very little forgetting. We did not detect any late-breaking transcriptional changes as sensitization memory was forgotten, nor was it possible to predict levels of forgetting from the state of single transcripts. We re-confirmed that a small set of transcripts remains persistently regulated after sensitization training.

This study completes a series conducted to trace the transcriptional correlates of sensitization memory as sensitization is encoded (Herdegen, Holmes, et al., 2014), maintained (Conte et al., 2017), forgotten (Perez et al., 2018), and then re-activated as a savings memory (Rosiles et al., 2020). Figure 5 integrates our findings from this paper with our previous work. In addition to characterizing the early and late waves of transcription produced by learning, our results shows that transcription and memory expression become dissociated after training: behavior can show strong expression of the memory with almost no detectable transcriptional changes, and the few transcriptional changes that persist do so relatively stably even as behavioral expression changes.

**Figure 5:**
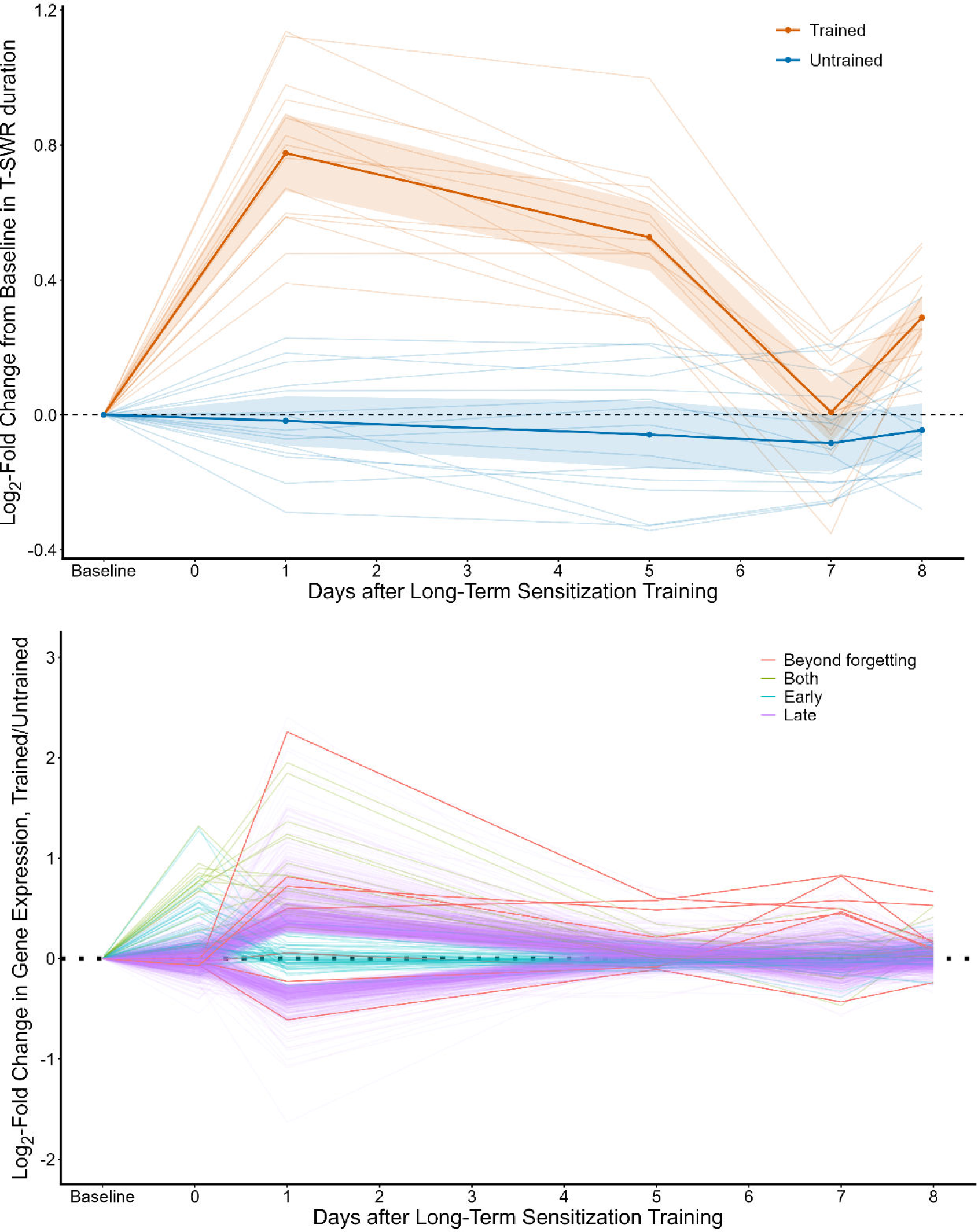
Comparison of the behavioral and transcriptional changes inducted by long-term sensitization training. A) The behavioral time course of long-term sensitization (replotted from Perez et al., 2018). Animals were given long-term sensitization training and also received a weak reminder shock after 7-day post-tests, revealing a latent sensitization memory. The plot shows T-SWR duration as the log-fold change from baseline on both the trained (red) and untrained (black) sides. B) The transcriptional time course of long-term sensitization (replotted from). This plot summarizes gene expression measured by microarray 1 hour (Herdegen et al., 2014), 1 day (Conte et al., 2017), 5 days (current study), 7 days (Perez et al., 2018), and 8 days but with a reminder shock applied after the 7-day post-tests (Rosiles et al., 2020). In total, 64 transcripts show clear regulation at 1 hour but not subsequently (“Early”, marked in teal); 1,237 show clear regulation at 1 day but not subsequently (“Late”, marked in purple), 17 are regulated at both 1 hour and 1 day but not subsequently (“Both”, brown), and 7 show regulation that at 1 and 7 days (“Beyond Forgetting”, red). Note that most transcriptional changes decay well before memory expression and that the return of memory expression after a reminder occurs in the absence of detectable changes in gene expression.

**Figure 6:**
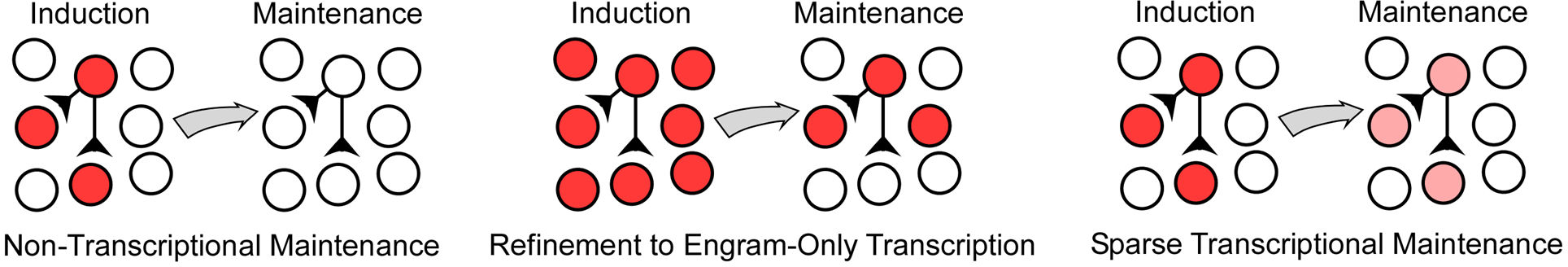
Mechanisms that would yield lower global changes in gene expression during memory maintenance. Left: One possibility is that memory maintenance does not require ongoing transcription, perhaps relying instead exclusively on post-transcriptional mechanisms. Middle: A second possibility is that the initial transcriptional response to sensitization is overly broad and is then refined only to neurons that will encode the memory. Right: A third possibility is that encoding neurons have a broad initial transcriptional response but require only a small subset of ongoing transcriptional changes to maintain the altered signaling that expresses sensitization.

The disconnect between transcription and memory expression is somewhat surprising. Specifically, sensitization has been proposed to activate self-sustaining transcriptional loops that then contribute to maintaining the expression of sensitization memory (Smolen et al., 2019). For example, cellular analogs of sensitization training induce CREB1 phosphorylation, increasing the binding of CREB1 to its own promotor, producing a long-term increase in the expression of CREB1 mRNA and protein (Liu, Shah, et al., 2011; Mohamed et al., 2005). This long-term increase in CREB1 transcription is essential for the long-term maintenance of cellular and synaptic correlates of long-term sensitization (Liu, Cleary, et al., 2011). Moreover, there is evidence that epigenetic changes are required to support the ongoing expression of sensitization (Pearce et al., 2017), and epigenetic marks are generally thought to alter neuronal phenotypes through alterered gene expression. Thus, recent reviews (Smolen et al., 2019) have posited that the maintenance of sensitization is at least partly due to ongoing transcriptional regulation. Our findings, in contrast, suggest that the vast majority of transcriptional changes induced after learning are transient, and that the few that do persist are not clearly linked to memory expression. What might explain this discrepancy between the hypothesis of transcription-mediated maintenance and the apparent discordance between transcription and memory expression?

One possibility is that we observe an early decay of transcription simply because there is a lag between the decay of transcription and the completion of forgetting, as the proteins produced by transcriptional regulation could extend behavioral expression for some time after gene expression returns to baseline conditions. While the notion of a lag between transcription and behavior makes sense, it would still be unclear why there are so few detectable transcription changes during savings memory (Rosiles et al., 2020). Nevertheless, we are now exploring more intensive training protocols that produce longer-lasting sensitization; this should provide a clearer test for maintenance-related transcription that is not muddied by dynamic behavioral expression.

Another possibility (Figure 5, Left) is that maintenance of sensitization is non-transcriptional. For example, cellular analogs of sensitization are accompanied by self-perpetuating conformational change in synaptically-expressed CPEB, a prion-like protein that regulates local translation (Miniaci et al., 2008; Si et al., 2003, 2010). This process is essential to early maintenance of synaptic correlates of long-term sensitization. Similarly, cellular analogs of sensitization also produce persistent-changes in post-translational processing of PKC (yielding PKM Apl III), and putative inhibitors of PKM have been reported to eliminate the expression of long-term sensitization in intact animals (Cai et al., 2011). Thus, there are already demonstrated maintenance mechanisms that operate solely on the protein level, and our data could be an indication these are the only key factors for sustaining the expression of sensitization. We cannot rule out this possibility, but we note a) that this makes it hard to understand why the initial transcriptional response is so broad, and b) long-term sensitization is not exclusively encoded via synaptic changes, long-term changes in excitability also contribute, and this at least suggests a cell-wide maintenance mechanism.

Another possibility is that the decline in transcriptional regulation we observe at the level of whole ganglia is due to a process of refinement. For example, the transcriptional response to sensitization training might be initially exuberant and then refined over time to just the subset of neurons that will encode the memory (Figure 5, Center). Or, perhaps, maintenance is transcriptionally sparse, requiring only a few ongoing changes after sensitization is consolidated (Figure 5, Right). Either of these scenarios would produce the ganglia-wide decline in transcriptional regulation we have observed, but this would belie continued transcriptional maintenance that has become harder to detect. We are now conducting screens with cell-type specificity, which should help clarify if we have missed ongoing but more-specific transcriptional changes.

Overall, the relationships between transcriptional, neuronal, and behavioral change remain surprisingly murky. While long-term memories require changes in neuronal gene expression at the time of learning, it is still difficult to provide a clear account of what, exactly, these diverse transcriptional changes accomplish, how long they last, and how/why forgetting can occur. Resolving these key issues will likely require time-course studies that can resolve transcription at the single-cell level with sufficient biological replicates to provide strong precision of measurement.

## Acknowledgements

This project is funded by NIMH grant 2R15MH107892-02

## Notes

https://osf.io/g3ueu

### Competing Interest Statement

The authors have declared no competing interest.

